# Simplified Molecular Classification of Lung Adenocarcinomas Based on *EGFR*, *KRAS*, and *TP53* Mutations

**DOI:** 10.1101/525949

**Authors:** Roberto Ruiz-Cordero, Junsheng Ma, Abha Khanna, Genevieve Lyons, Waree Rinsurongkawong, Roland Bassett, Ming Guo, Mark J. Routbort, Jianjun Zhang, Ferdinandos Skoulidis, John Heymach, Emily B. Roarty, Zhenya Tang, L. Jeffrey Medeiros, Keyur P. Patel, Rajyalakshmi Luthra, Sinchita Roy Chowdhuri

## Abstract

**Introduction:** Gene expression profiling has consistently identified three molecular subtypes of lung adenocarcinoma that have prognostic implications. To facilitate stratification of patients with this disease into similar molecular subtypes, we developed and validated a simple, mutually exclusive classification.

**Methods:** Mutational status of *EGFR*, *KRAS*, and *TP53* was used to define six mutually exclusive molecular subtypes. A development cohort of 283 cytology specimens of lung adenocarcinoma was used to evaluate the associations between the proposed classification and clinicopathologic variables including demographic characteristics, smoking history, fluorescence in situ hybridization and molecular results. For validation and prognostic assessment, 63 of the 283 cytology specimens with available survival data were combined with a separate cohort of 428 surgical pathology specimens of lung adenocarcinoma.

**Results:** The proposed classification yielded significant associations between these molecular subtypes and clinical and prognostic features. We found better overall survival in patients who underwent surgery and had tumors enriched for *EGFR* mutations. Worse overall survival was associated with older age, stage IV disease, and tumors with comutations in *KRAS* and *TP53*. Interestingly, neither chemotherapy nor radiation therapy showed benefit to overall survival.

**Conclusions:** The mutational status of *EGFR*, *KRAS*, and *TP53* can be used to easily classify lung adenocarcinoma patients into six subtypes that show a relationship with prognosis, especially in patients who underwent surgery, and these subtypes are similar to classifications based on more complex genomic methods reported previously.

## Introduction

Lung cancer is the main cause of cancer-related mortality in both men and women^1-3^. Lung adenocarcinoma accounts for approximately 40% of lung cancer cases^4-6^. Gene expression profiling (GEP) of lung adenocarcinomas has consistently identified three molecular subtypes with prognostic implications^7-14^. The initial molecular classification of lung adenocarcinomas included the bronchoid, magnoid, and squamoid subtypes^11,15^. However, after comprehensive molecular profiling of a cohort of lung adenocarcinomas, The Cancer Genome Atlas Research Network proposed an updated nomenclature for this molecular classification that encompasses previous histopathologic, anatomic, and mutational classifications^13^. This system re-designated the initial subtypes as the terminal respiratory unit, proximal-proliferative, and proximal-inflammatory subtypes, respectively^13^.

Tumors with acinar, papillary, or lepidic histomorphology and mutations or copy number alterations in *EGFR*, presenting most often in women who have never smoked, predominantly cluster in the terminal respiratory unit subtype. Tumors in the proximal-proliferative subtype have variable histology and commonly display mutations and copy number alterations in *KRAS* and *STK11*. In contrast, lung adenocarcinomas with primarily solid architecture and enrichment for *TP53* and *NF1* mutations and *p16* methylation typically cluster in the proximal-inflammatory subtype^13,15^.

While molecular subtypes of lung adenocarcinoma have been associated with significant differences in prognosis, routine GEP in the clinical setting has been limited by cost, complexity, and increased turnaround time^16^. These limitations have led to the development of simplified prognostic models based on the expression of selected genes^10,16^. However, many of these genes, such as *PTK7, CIT, SCNN1A, PTGES, ERO1A, ZWINT, DUSP6, MMD, STAT1, ERBB3*, and *LCK*, are not tested routinely in the clinical laboratory.

To fill this need, we developed a simplified molecular subtype classification based on the mutational status of only *EGFR, KRAS*, and *TP53* to facilitate categorization of patients’ lung adenocarcinomas into molecular subtypes with relevant prognostic information.

## Materials and Methods

### Patient selection for development cohort

We retrospectively reviewed our institutional database for patients treated between May 1, 2010, and October 31, 2015, to identify cytologic specimens of patients with lung adenocarcinoma. Patients with TTF1-negative non–small cell lung cancer, small cell lung cancer, large cell carcinoma, squamous carcinoma, and poorly differentiated carcinoma not otherwise specified were excluded. We reviewed the patients’ medical records for demographic characteristics, clinical information, fluorescence in situ hybridization (FISH) results for *ALK*, *ROS1*, *MET*, and/or *RET*, and mutation profiling data derived by next-generation sequencing (NGS) and polymerase chain reaction (PCR)-based methods (i.e. Sanger sequencing or pyrosequencing). PCR-based methods were restricted to analysis of only *EGFR*, *KRAS*, and *BRAF* hotspots.

### Patient selection for validation cohort

Patients from our institution’s Genomic Marker-Guided Therapy Initiative (GEMINI) project database were selected as a validation cohort. This group included patients who underwent computerized tomography–guided transthoracic core-needle biopsy for diagnosis and/or staging of lung adenocarcinomas as well as patients who underwent surgery to resect lung adenocarcinoma between November 1, 2009, and October 31, 2016. Age, sex, race/ethnicity, smoking status, NGS mutation data, survival status, and treatment information were included in the analysis. To avoid Simpson’s paradox^17^, we combined this cohort with a subset of cytology cases from the development cohort whose medical record numbers matched to those of records in the GEMINI database and who had available survival information and treatment data.

### Mutational analysis

NGS was performed on cytology smears or formalin-fixed paraffin-embedded tissue (cytology cell blocks or core biopsy tissue blocks) using the Ion Torrent or Ion Proton (Thermo Fisher Scientific) sequencers in our College of American Pathologists– accredited, Clinical Laboratory Improvement Amendments–certified laboratory. Multiple NGS panels were developed, validated, and implemented in our laboratory during the study period (2009-2016), including an initial hotspot panel of 46 cancer-related genes^18^, an updated 50-gene hotspot panel, a 126-gene panel, and a panel of 409 cancer-associated genes^19^. The cytology specimens were appropriately validated^20^. All these panels include several amplicons targeting known hotspots in exons of *EGFR*, *KRAS*, and *TP53*.

### Simplified classification of molecular subtypes

We stratified cases from our development and validation cohorts by creating a classification system using the mutational status of *EGFR*, *KRAS*, and *TP53*, forming mutually exclusive groups. Cases that harbored mutations in *EGFR* only or mutations in *EGFR* and genes other than *KRAS* and *TP53* were classified as the simplified terminal respiratory unit (sTRU) subtype. Cases with *KRAS* mutations only or mutations in *KRAS* and genes other than *EGFR* and *TP53* were classified as the simplified proximal-proliferative (sPP) subtype. Cases with only *TP53* mutations or mutations in *TP53* and genes other than *EGFR* and *KRAS* were classified as the simplified proximal-inflammatory (sPI) subtype. Also, cases with co-mutations in *KRAS* and *TP53* (*KRAS/TP53* subtype) or *EGFR* and *TP53* (*EGFR/TP53* subtype) were grouped separately. Cases with mutations in genes other than *EGFR*, *KRAS*, and *TP53* were classified as the non-TRUPPPI subtype, and a few cases that lacked mutations in any of the genes detected by our NGS panels were placed in a “no-mutation” subtype.

## Statistical Methods

### Development cohort

Categorical variables were summarized by frequencies and percentages, and continuous variables were summarized using means, standard deviations, medians, and ranges. Fisher exact test or its generalization for categorical variables was used to compare categorical variables between molecular subtypes; in addition, Monte Carlo simulation approach was used when computational issues were encountered. Patients with indeterminate FISH results or unknown aneuploidy status were excluded from the Fisher exact tests. The Wilcoxon rank sum test or Kruskal-Wallis rank-sum test was used to compare continuous variables between molecular subtypes.

### Validation cohort

Associations between variables and subtypes were assessed as described for the development cohort. The outcome variable of overall survival (OS) time was computed from the date of initial diagnosis to the last follow-up date or death date. For the subset of patients who had surgery, separate analyses were performed of OS from the date of surgery. Cox proportional hazards models were used to evaluate associations of variables with survival outcomes, and Firth penalized Cox regression models were fitted for covariates with zero count of events. In multivariate Cox regression analyses, we included covariates that had p values less than 0.25 in univariate Cox regression models. Treatment variables (surgery, radiation, and chemotherapy) were handled as time-varying covariates. The Kaplan-Meier method was used to estimate survival distributions, and the log-rank test was used for comparisons between survival distributions. All statistical analyses were performed using R version 3.3.11^21^ and SAS version 9.4. All statistical tests used a significance level of 5%, and no adjustments for multiple testing were made.

## Results

### Development cohort

We collected a development cohort of 283 consecutive cytology samples from patients with lung adenocarcinoma. The samples were acquired via endobronchial ultrasound–guided FNA (64.7%, n=183), thoracentesis and paracentesis (16.6%, n=47 [46 pleural samples]), computed tomography–guided FNA (13.4%, n=38), and ultrasound-guided FNA (5.3%, n=15); most samples were cell block preparations (97.8%, n=277). Metastases accounted for 82.7% (n=234) of cases, and the majority were to lymph nodes (66.7%, n=156), followed by pleural fluid (20.1%, n=47), soft tissue (3.8%, n=9), bones (3.4%, n=8), adrenal glands (3.0%, n=7 [5 on the left]), liver (2.1%, n=5), and other sites (0.9%, n=2). The relevant demographic characteristics and clinicopathologic data for the development cohort are summarized in Table 1. The cohort was composed primarily of older individuals with a median age of 65.4 years (range: 27.5-90.2 years) and 151 (53.4%) women. Most patients were white, current or former smokers, and had stage IV disease at the time of data collection. All cases underwent FISH testing for *ALK*, *ROS1*, *MET*, and/or *RET.* The FISH results were negative in 250 (88.3%) cases. The rest of the cases were positive for rearrangements or amplification of *ALK* (5.7%, n=16), *MET* (1.8%, n=5), *RET* (0.7%, n=2), or *ROS1* (0.3%, n=1), or were indeterminate (3.2%, n=9). Aneuploidy, defined as an increase or decrease in the number of fluorescent signals observed in a cell, was present in 193 (68.2%) cases, not present in 60 (21.2%), indeterminate in 27 (9.5%), and not assessed in three (1.1%) cases.

**Table 1.** Demographic and clinicopathologic characteristics in the development cohort.

| Variable | Value |
| --- | --- |
| **Sex, n (%)** |  |
| Male | 132 (46.6) |
| Female | 151 (53.4) |
| **Age, median (range)** | 65.4 y (27.5-90.2 y) |
| **Race/ethnicity, n (%)** |  |
| Caucasian | 215 (76.0) |
| Asian | 27 (9.5) |
| Black | 19 (6.7) |
| Hispanic | 21 (7.4) |
| Unknown | 1 (0.3) |
| **Smoking status, n (%)** |  |
| Never-smoker | 64 (22.6) |
| Former smoker | 136 (48.1) |
| Current smoker | 82 (29.0) |
| Unknown | 1 (0.3) |
| **Vital status, n (%)** |  |
| Alive | 172 (60.8) |
| Deceased | 111 (39.2) |
| **Molecular platform, n (%)** |  |
| NGS | 218 (77) |
| PCR* | 55 (19.5) |
| Not done | 10 (3.5) |
| **Molecular subtype, n (%) (n=233)** |  |
| sTRU – *EGFR* (%) | 34 (14.6) |
| sPP – *KRAS* (%) | 43 (18.5) |
| sPI – *TP53* (%) | 46 (19.7) |
| Co-mutation | 60 (25.8) |
| *EGFR/TP53* | 26 (11.2) |
| *KRAS/ TP53* | 34 (14.6) |
| Non-TRUPPPI | 21 (9.0) |
| No mutations detected by NGS | 29 (12.4) |
| **FISH results, n (%)** |  |
| Negative | 250 (88.3) |
| Positive | 24 (8.5) |
| *ALK* | 16 (5.7) |
| *ROS1* | 1 (0.4) |
| *RET* | 2 (0.7) |
| *MET* | 5 (1.8) |
| Aneuploidy | 193 (68.2) |

\*Sanger sequencing or pyrosequencing
NGS, next-generation sequencing; PCR, polymerase chain reaction; sTRU, simplified terminal respiratory unit; sPP, simplified proximal-proliferative; sPI, simplified proximal-inflammatory; non-TRUPPPI, mutations in genes other than *EGFR*, *KRAS*, and *TP53*; FISH, fluorescence in situ hybridization

Mutational analysis, including NGS and PCR-based methods, was performed in 273 (96.5%) cases. NGS was performed in 77% (n=218) of the specimens, yielding positive mutations in 188 (86.2%) cases. PCR-based testing was performed in 55 (20%) cases, yielding positive mutations in 15 (27.3%) (9 in *EGFR*, 6 in *KRAS*, and 0 in *BRAF*). Of the cases with PCR-based testing, two cases had inadequate DNA material, and 38 cases were negative for single-gene testing. These 40 cases as well as 10 cases in which mutational analysis was not performed were excluded from further analysis, leaving 233 cases. Mutations were most frequent in *TP53* (46.0%, n=107), *KRAS* (33.5%, n=78), and *EGFR* (26.2%, n=61). According to our proposed classification, 34 (14.6%) cases were classified as sTRU, 43 (18.5%) as sPP, and 46 (19.7%) as sPI. Cases with co-mutations included 26 (11.2%) with *EGFR/TP53* and 34 (14.6%) with *KRAS/TP53*. There were 21 (9%) cases with mutations in genes other than *EGFR*, *KRAS*, and *TP53* (non-TRUPPPI subtype) and 29 (12.4%) cases with no mutations detected.

The simplified molecular subtypes were statistically significantly associated with age, race/ethnicity, smoking status, and aneuploidy (Table 2). To identify further associations, we compared variables between patients within a given molecular subtype and the remaining patients. The sTRU subtype was associated with Asian race/ethnicity (23.5% vs. 7.6%, p=0.027) and never-smoker status (52.9% vs. 18.7%, p<0.001). The sPP subtype was associated with white race/ethnicity (86.0% vs. 73.0%, p=0.042). The sPI subtype was associated with male sex (63.0% vs. 41.2%, p=0.008). The *EGFR/TP53* subtype was associated with younger age (mean age 56.9 vs. 65.8 years, p<0.001), Asian (19.2% vs. 8.7%) and Hispanic race/ethnicity (19.2% vs. 6.3%, p=0.026), never-smoker status (46.2% vs. 20.9%, p=0.016), and lack of aneuploidy (60.0% vs. 80.1%, p=0.038). The *KRAS/TP53* subtype was associated with current smoking (55.9% vs. 21.7%, p<0.001). The non-TRUPPPI subtype was not associated with any of the covariates, and the no-mutation subgroup was associated with never-smoker status (41.4% vs. 21.2%, p=0.045) and aneuploidy (95.8% vs. 75.3%, p=0.019).

**Table 2.**
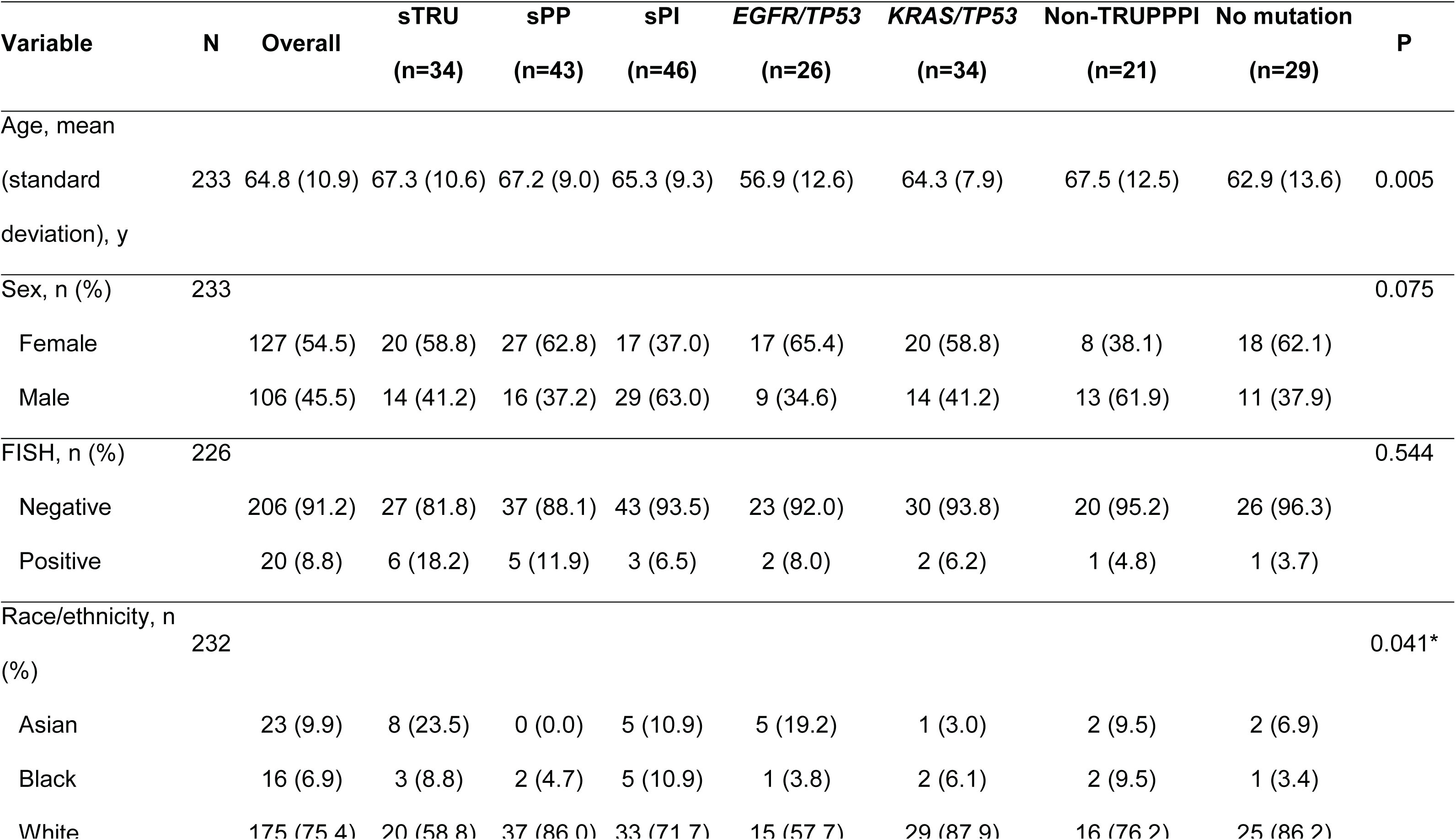
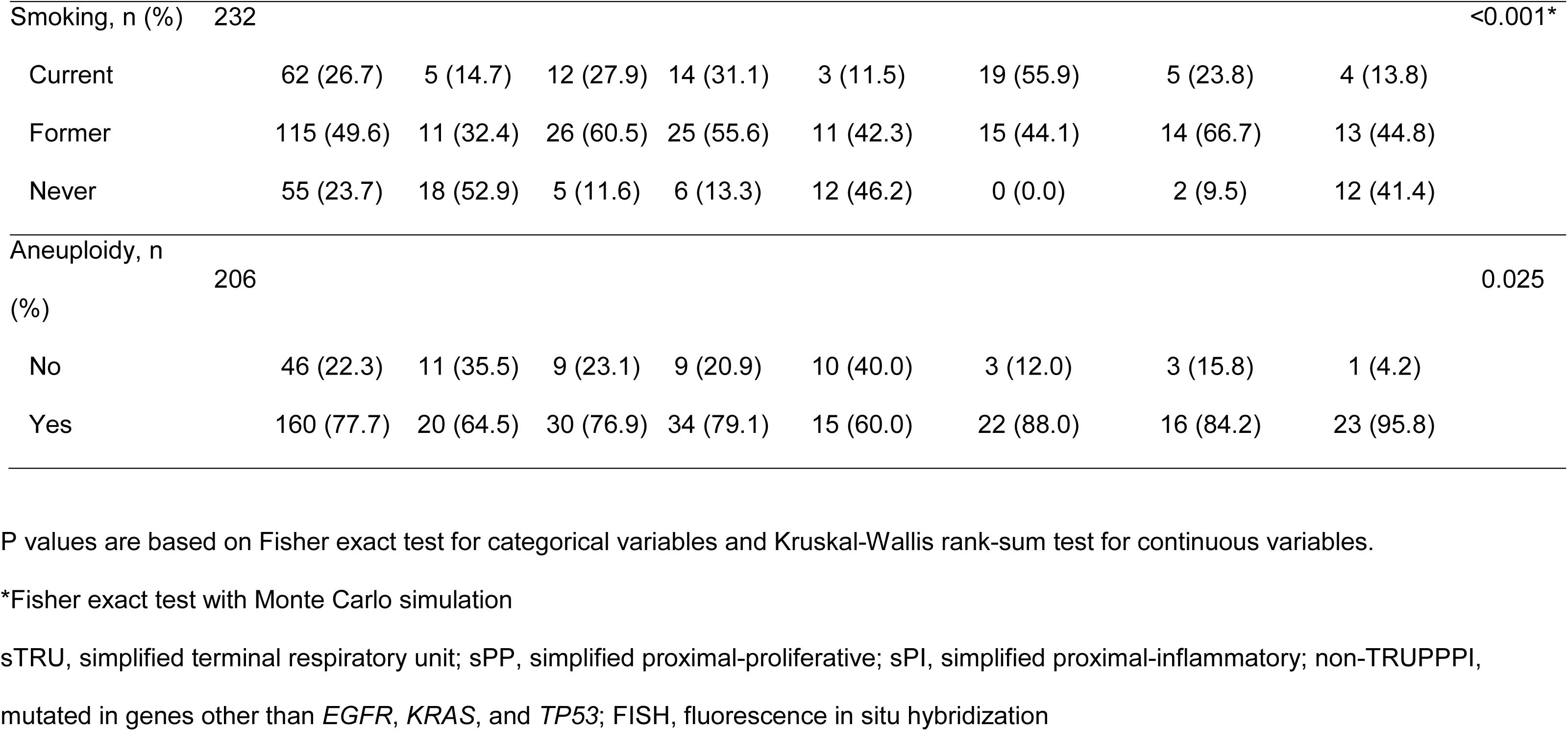
Associations between molecular subtypes and clinicopathologic variables in the development cohort.

### Validation cohort

To validate these findings and determine the impact of our subtypes on prognosis, we used a validation cohort (n=428) composed of core-needle biopsy samples or resection specimens from lung adenocarcinoma patients with available data on treatment and follow-up. Histomorphologic subtypes (e.g., mucinous, lepidic, acinar, and solid) were reported in 28.3% (n=121) of the pathology reports. The mutational data for this cohort were based only on NGS because all three target genes were not assessed in cases where PCR-based single-gene analysis was performed. Also, we included the 63 patients from the cytology cohort in the GEMINI database with treatment and follow-up data available. NGS results were available for 85.7% (n=54) of these cases.

#### Mutational profiling of lung adenocarcinoma patients in the validation cohort

Sequencing data were available for 484 (98.6%) patients in the combined validation cohort. NGS and PCR analyses yielded a total of 835 mutations/variants in 421 patients (87.0%). The median tumor percentage was 40% (range: 20 to 95 % tumor cells). Most of the genomic alterations were missense mutations (75%, n=618), followed by in-frame deletions (7%, n=58), nonsense (6.6%, n=55) and frameshift (5%, n=40) mutations, duplications (2.1%, n=18), complex mutations/indels (1.8%, n=15), splice mutations (1.4%, n=12), and gene amplifications (1.1%, n=10). Transversions included G>T (27%, n=222), T>G (7%, n=60), C>A (1.5%, n=13), and A>C (1%, n=8), and transitions included G>A (13%, n=106), C>T (12%, n=99), A>G (4%, n=34), and T>C (1%, n=9). The most common protein alterations were KRAS-G12C (n=58), EGFR-L858R (n=51), EGFR-E746_A750del (n=45), KRAS-G12V (n=36), KRAS-G12D (n=27), and EGFR-T790M (n=27). The mutational data for all 491 cases in the validation cohort, stratified by simplified molecular subtype, are summarized in Figure 1.

**Figure 1.**
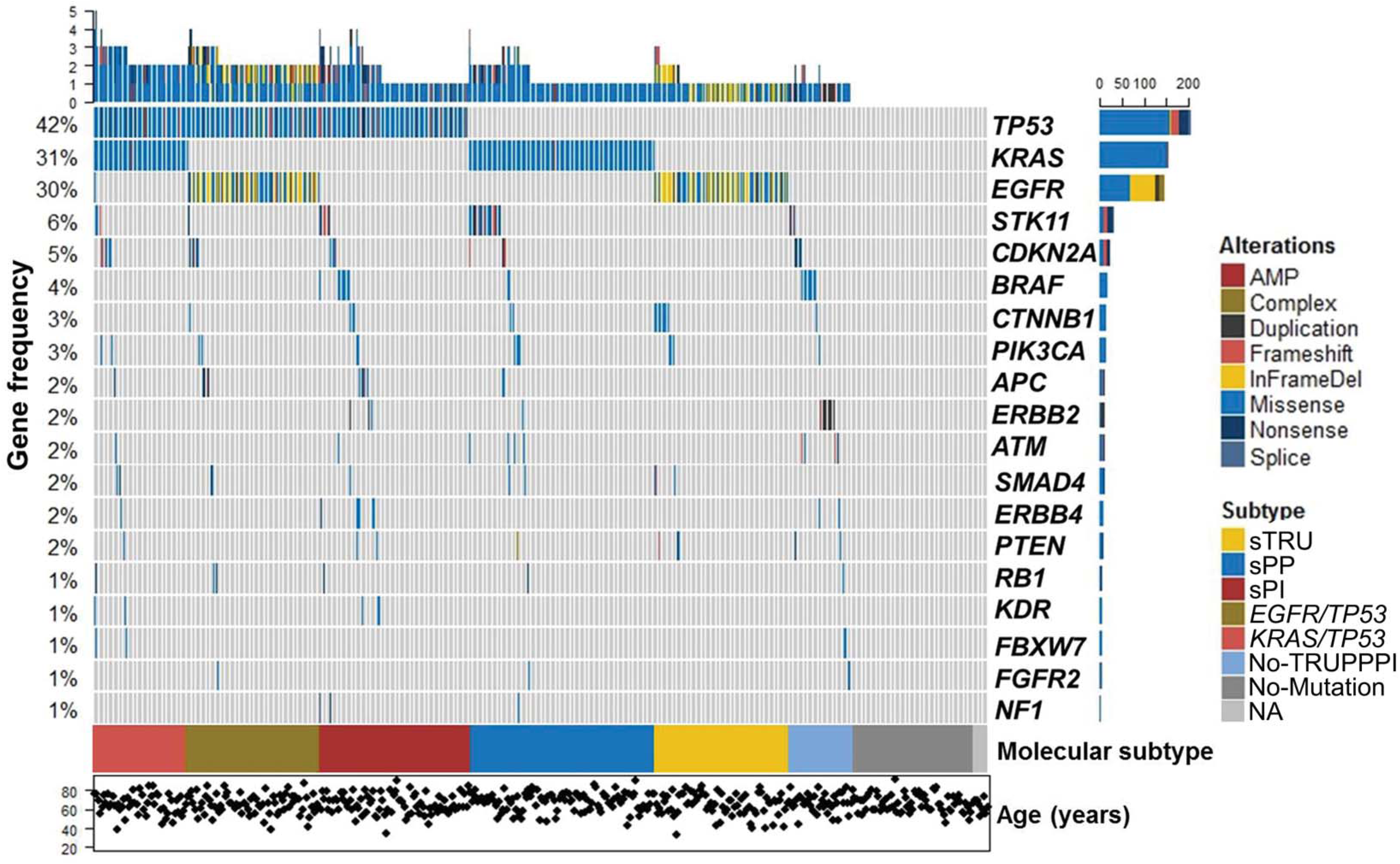
Oncoprint plot of the validation cohort. The mutational profile of the 491 patients in the validation cohort including the 19 most commonly mutated genes is shown. Each column represents a patient. The upper histogram highlights the number of genes mutated in each case. Mutated genes are in descending order of frequency, and their mutation frequency is shown on the y axis. Below the columns, the color map indicates the simplified molecular subtypes. At the bottom, the dot plot shows the age of each patient. The horizontal bars adjacent to the genes illustrate the number and type of genetic alterations. AMP, amplification; InFrameDel, in-frame deletion; NA, sequencing not available.

#### Clinical and histomorphologic associations according to simplified molecular subtypes

The simplified molecular subtypes were significantly associated with age, race/ethnicity, sex, smoking status, stage, histomorphology and FISH results (Table 3). As in the development cohort, variables were compared between patients within a given molecular subtype and the remaining patients. The sTRU subtype was associated with slightly older age (mean age 66.4 vs. 63.2 years, p=0.015), never-smoker status (62.2% vs. 26.3%, p<0.001) Asian race/ethnicity (17.6% vs. 6.6%, p=0.022), metastatic tumors (61.5% vs. 41.7%, p=0.013), non-mucinous (95.5% vs. 70.7%, p=0.014) and lepidic histology (63.6% vs. 36.4%, p=0.030), and stage IV disease (55.4% vs. 41.6%, p=0.007). The sPP subtype was associated with slightly older age (65.9 vs. 63.1 years, p=0.026), lower likelihood of never-smoker status (10.8% vs. 37.4%, p<0.001), black race/ethnicity (10.8% vs 6.5%, p=0.033), tumors with mucinous (42.9% vs. 17.4%, p=0.005) and non-acinar (80.0% vs. 58.1%, p=0.035) histology, non-metastatic tumors (69.1% vs 50.9%, p=0.016), negativity for *ALK/ROS1/MET/RET* abnormalities (96.8% vs 86.6%, p=0.003), and stage II disease (18.6% vs. 7.1%, p<0.001). The sPI molecular subtype was associated with male sex (55.4% vs. 39.4%, p=0.01), lower likelihood of never-smoker status (16.9% vs. 34.9%, p=0.003), and stage III disease (32.5% vs. 20.2%, p=0.047). Like the sTRU subtype, the *EGFR*/*TP53* subtype was associated with younger age (mean age 59.1 vs. 64.5 years, p<0.001), Asian race/ethnicity (18.9% vs. 6.3%, p<0.001), never-smoker status (55.4% vs. 27.6%, p<0.001), and smaller tumors (mean tumor size 3.27 cm vs. 3.96 cm, p=0.042). In contrast, the *KRAS*/*TP53* subtype was associated with Hispanic race/ethnicity (12.0% vs. 5.8%, p=0.035), lower likelihood of never-smoker status (6.0% vs. 34.8%, p<0.001), and solid (45.5% vs. 11.8%, p=0.011) and non-lepidic (90.9% vs. 55.5%, p=0.026) histology. The non-TRUPPPI subtype was associated with mucinous histology (62.5% vs. 22.1%, p=0.022), whereas the no-mutation subtype was associated with never-smoker status (46.0% vs. 29.7%, p=0.035) and acinar histology (57.1% vs. 31.0%, p=0.043). No significant associations were identified between subtype and alcohol intake.

**Table 3.**
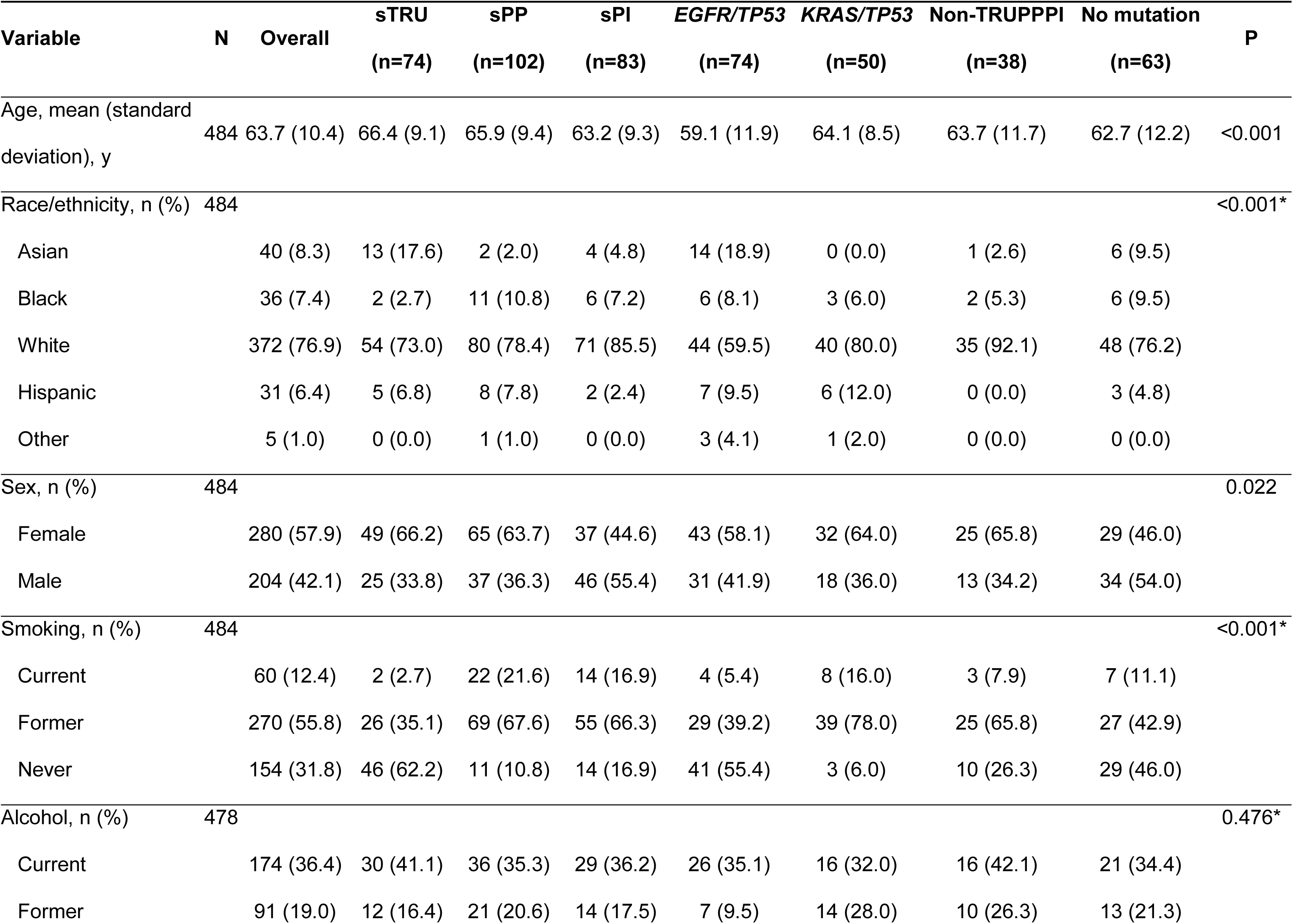
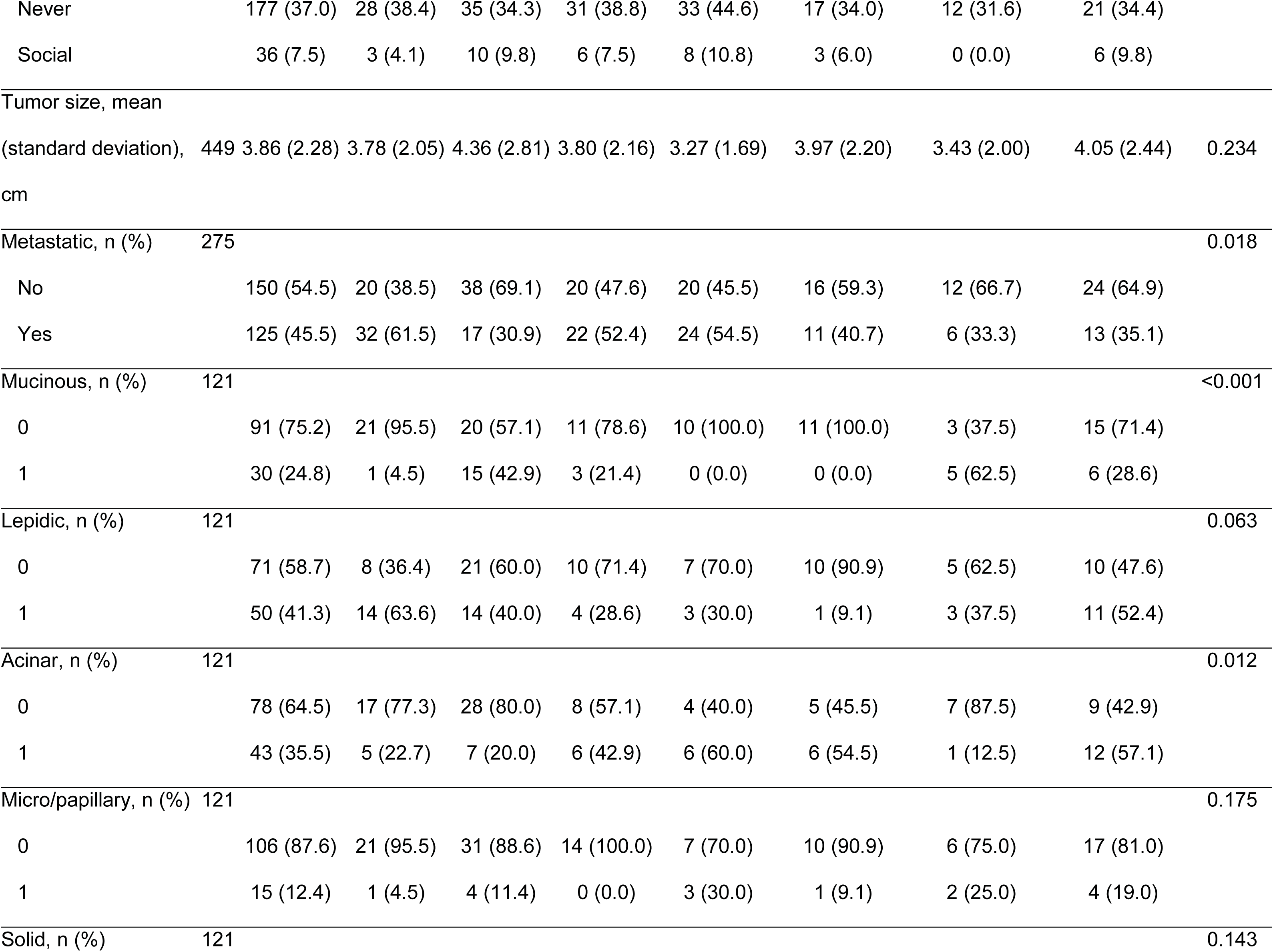
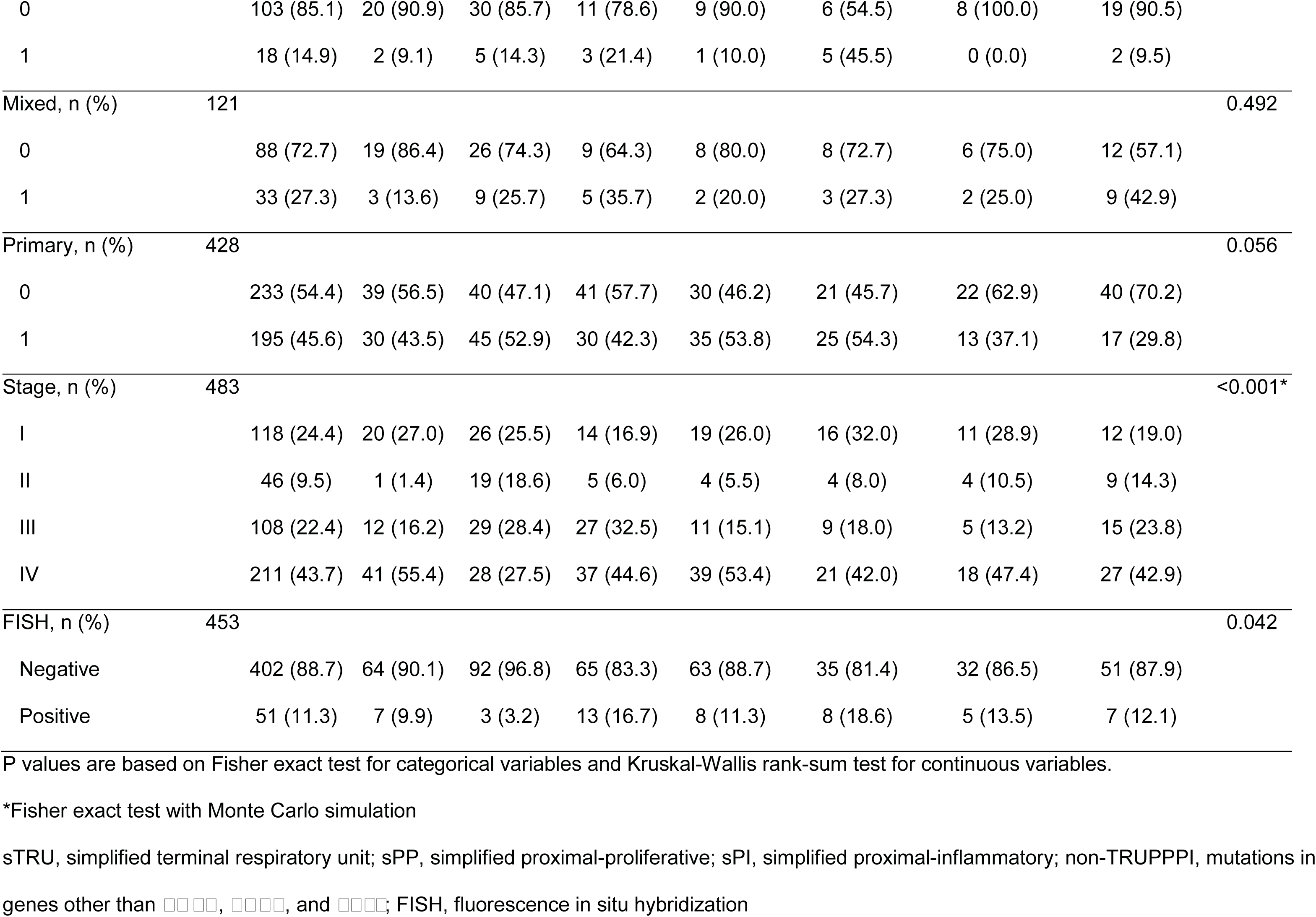
Associations between molecular subtypes and clinicopathologic variables in the validation cohort.

#### Prognostic associations according to simplified molecular subtype classification

We assessed overall survival in the validation cohort as previously described. The median follow-up time was 1.87 years (interquartile range: 0.9-3.5 years). The median survival time was 5.93 (95% CI: 4.57-not reached) years. We fitted a multivariate Cox proportional hazards regression model to assess for associations between OS and the covariates of age, sex, alcohol intake, stage, molecular subtype, and treatment (surgery, radiation, and/or chemotherapy), selected on the basis of univariate analyses with a cutoff p value of 0.25. We observed that patients with older age (hazard ratio [HR]=1.03, 95% CI: 1.01-1.05) and stage IV disease (HR=6.51, 95% CI: 2.49-17.04, p=0.006) had worse OS. Although the difference was not statistically significant, patients in the sTRU subtype had better OS (HR=0.42, 95% CI: 0.18-1.00, p=0.051) whereas patients in the *KRAS*/*TP53* subtype had worse OS (HR=2.15, 95% CI: 1.02-4.53, p=0.043) than those in the other subtypes (Figure 2A & B). Patients who underwent surgical resection (HR=0.33, 95%CI: 0.18-0.60, p<0.001) had better OS than those who did not have surgery. Figure 3A shows significant differences in OS within this subset of patients. We observed statistically significant differences in OS between the sTRU, sPP, and sPI subtypes (Figure 3B), however, differences between these subtypes when categorized in early stages I and II or late stages III and IV did not reach statistical significance (not shown). Interestingly, when compared with patients who underwent surgery, no significant differences in OS were observed in patients who received chemotherapy, and OS was significantly worse in those who received radiation therapy, regardless of molecular subtype (HR=1.87, 95% CI: 1.18-2.96, p=0.007). Notably, OS did not significantly differ between the sTRU and *EGFR/TP53* subtypes (log-rank test p=0.84), either in all patients (not shown) or in the patients who underwent surgery (Figure 3C), suggesting that these subtypes could represent a single group. Conversely, the *KRAS/TP53* subtype showed the poorest OS both among all patients (HR=2.15, 95% CI: 1.02-4.53, p=0.043) (Figure 2B) and among those who underwent surgery (HR=1.935, 95% CI: 0.923-4.058) (Figure 3D).

**Figure 2.**
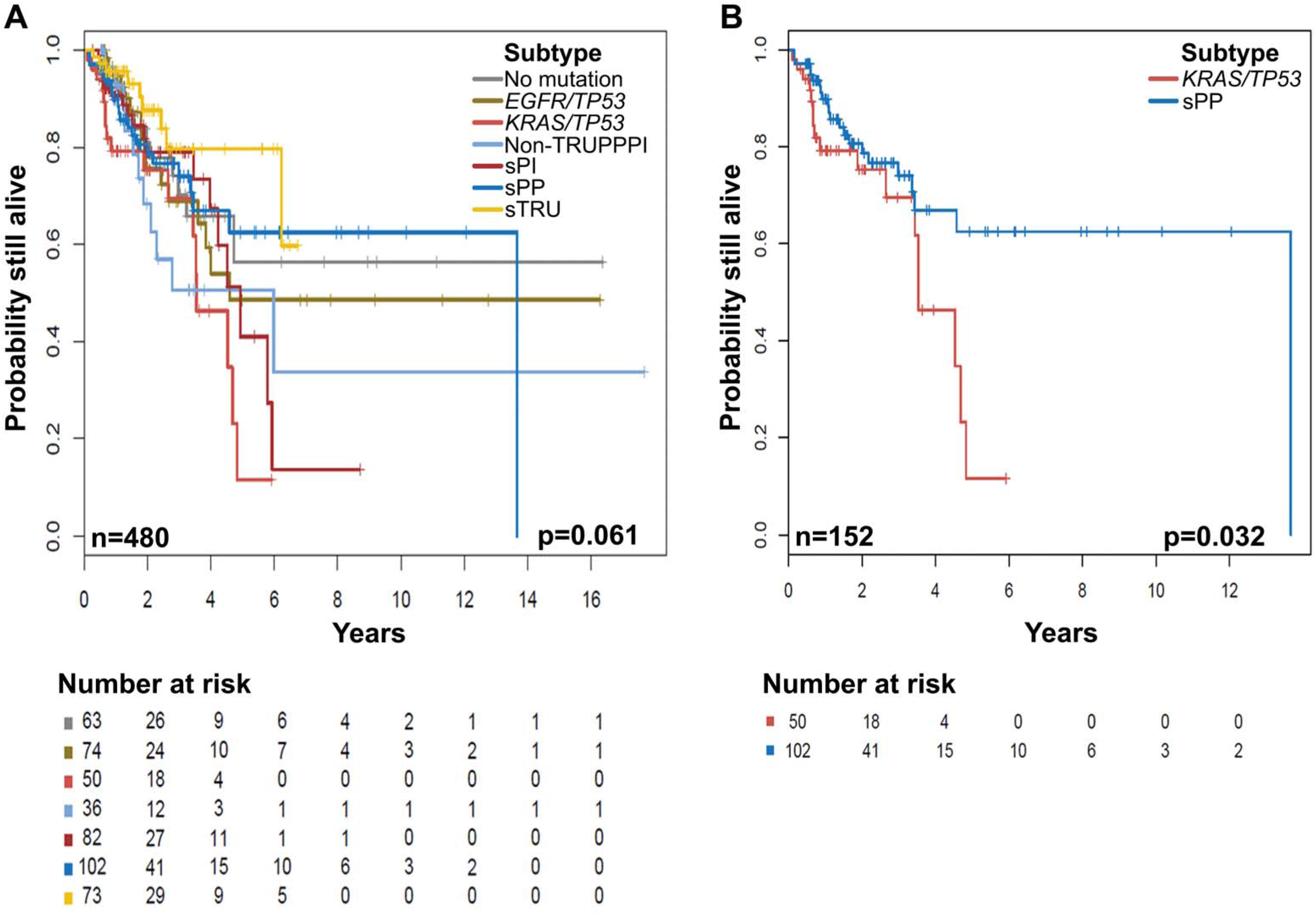
Overall survival (OS) stratified by simplified molecular subtypes. Kaplan-Meier plots show the prognostic significance of the simplified molecular subtypes. **A)** OS differs between the simplified molecular subtypes. **B)** OS differs significantly between the sPP and *KRAS/TP53* subtypes. The number of patients and the log-rank test p value are shown at the bottom of each plot.

**Figure 3.**
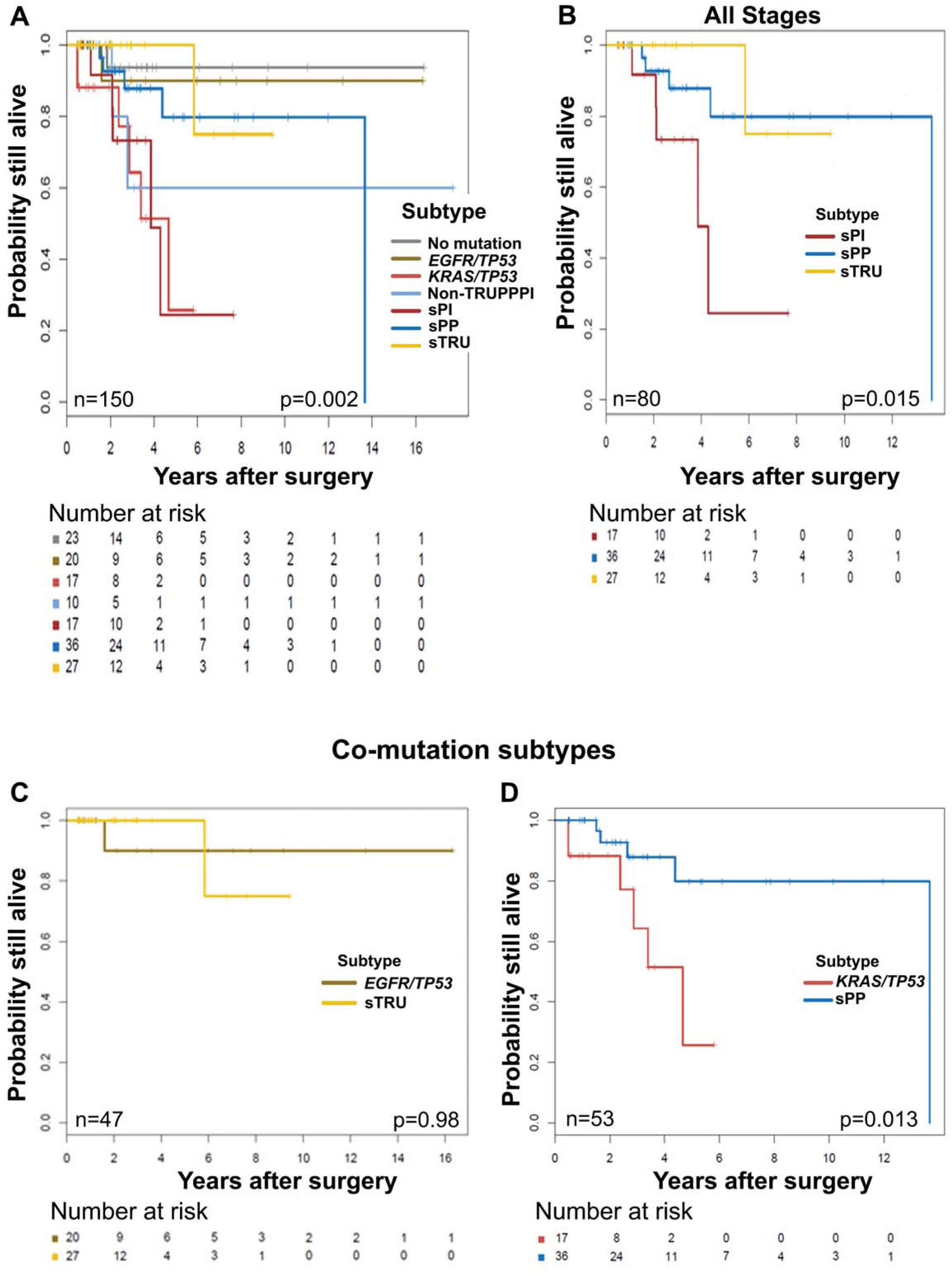
Overall survival (OS) in patients undergoing surgery stratified by simplified molecular subtypes. Kaplan-Meier plots show the prognostic significance of the simplified molecular subtypes in patients who underwent surgery. **A)** OS significantly differs between the simplified molecular subtypes. **B)** sPI shows worse OS than sTRU and sPP when including all tumor stages. **C)** OS does not significantly differ between the sTRU and *EGFR/TP53* subtypes. **D)** OS does significantly differ between the sPP and *KRAS/TP53* subtypes. The number of patients and the log-rank test p value are shown at the bottom of each plot.

## Discussion

In this study, we show that the mutational status of three commonly mutated genes can be utilized to create a simplified, mutually exclusive molecular subtype classification of lung adenocarcinomas based on molecular subtypes previously identified using GEP or larger gene mutation panels and that this simplified classification shows a relationship with prognosis, especially in patients who have undergone surgery.

The simplified classification showed high concordance with most previously reported associations, but there were some notable differences. For example, most patients in the sTRU subtype had advanced-stage disease. Among advanced-stage cases, however, those with sTRU and *EGFR/TP53* subtypes had a better prognosis, perhaps a reflection of our referral patient population, whose disease often has not responded to first-line therapy and who present with high-grade, advanced-stage tumors. As expected, the sTRU subtype was also enriched for Asian patients with better prognosis and lepidic histology^11,13,22^. Our results also suggest that adenocarcinoma histologic types do not correlate with stage, in keeping with previous findings^11^. However, we observed significant associations between some of the simplified molecular subtypes and morphology. As suggested by Nakaoku et al. and others^23,24^, the PP subtype as well as our sPP subtype are associated with mucinous histology. While the sTRU subtype was not associated with lepidic histology as the TRU is^25^, the sTRU however, was associated with non-mucinous tumors. *KRAS/TP53*-mutated tumors more often had solid histology; similarly, a study by Rekhtman et al.^24^ found a significant association between a subset of *KRAS*-mutated tumors and solid histology; however, they did not test for *TP53* mutations. Our observations suggest that this association could be unique to *KRAS/TP53* co-mutated tumors.

Interestingly, acinar histology was common in tumors that did not show mutations in this study. Since our NGS panels were developed to target specific exons and did not provide whole-exome/genome results, the no-mutation group could harbor infrequent intronic or exonic mutations/polymorphisms in *EGFR*, *KRAS*, *TP53*, and/or other genes. Whereas genomic alterations have been found in all tumors tested by various groups^26,27^, tumors with rare alterations of currently unknown significance could represent unique subtypes where oncogenesis is not driven by common mutations in known genes and genetic pathways^28^.

No major differences were observed between the sTRU and *EGFR/TP53* subtypes. We therefore suggest that *EGRF/TP53* cases can be combined with sTRU cases. However, the sPP and *KRAS/TP53* subtypes must be clearly distinguished since the latter appears to confer the poorest OS. A report from our group demonstrated different subtypes within *KRAS*-mutated cases and further supports that *KRAS/TP53* co-mutation portends the worst prognosis^29^.

Our findings concord and confirm with previous findings that radiation^30^ and chemoradiotherapy carry a worse OS with more toxicity and a higher rate of death during treatment, particularly in older patients^31^. Our work thus builds upon recent evidence suggesting radiation therapy be reconsidered in patients with lung adenocarcinoma.

In keeping with the recent recommendations by the updated molecular testing guidelines for the selection of lung cancer patients for targeted therapy, our results provide additional support for the use of cytology specimens as a valuable sample source for molecular testing in patients with lung adenocarcinoma. NGS further enables testing of FNA material and helps avoid the potential risks associated with surgical biopsies^32-39^.

Others have shown that GEP using microarray technology reliably estimates prognosis^10,16^, but the use of microarrays in the clinical setting is limited by the large number of analyzed genes, complex methods, independent validation of the results, low inter-laboratory reproducibility, high cost, long turnaround time, and the need for fresh or frozen tissue^40^. By creating mutually exclusive groups based on easily accessible data, such as *EGFR*, *KRAS*, and *TP53* status, the classification of lung adenocarcinomas into prognostic molecular subtypes could become readily available in routine clinical practice. While oversimplification is a potential limitation of the classification proposed, we believe this simplified classification provides useful prognostic information while retaining the updated proposed nomenclature (i.e., TRU, PP, and PI). This simplified approach will make it easier for molecular genetics laboratories and clinicians to accurately classify patients and will help maintain consistency across different molecular laboratories employing NGS platforms for genomic analysis. Because of the increasing demand for multigene testing over single-gene tests^39^ and because most, available NGS panels testing lung adenocarcinoma samples contain these three key genes, we suggest that this simplified classification be used primarily for results obtained via NGS.

In summary, using mutational data for *EGFR*, *KRAS*, and *TP53*, we have defined prognostic groups similar to those previously identified by more complex genomic methods in patients with lung adenocarcinomas.

## Acknowledgments

Authors would like to thank Ms. Sarah J. Bronson for her editorial support.

